# Hypoxia in the grape berry linked to mesocarp cell death: the role of seed respiration and lenticels on the berry pedicel

**DOI:** 10.1101/209890

**Authors:** Zeyu Xiao, Suzy Rogiers, Victor Sadras, Stephen D. Tyerman

## Abstract

Mesocarp cell death (CD) during ripening is common in berries of seeded *Vitis vinifera* L wine cultivars. We examined if hypoxia within berries is linked to CD. Internal oxygen concentration ([O_2_]) across the mesocarp was measured in berries from Chardonnay and Shiraz, both seeded, and Ruby Seedless, using an oxygen micro-sensor. Steep [O_2_] gradients were observed across the skin and [O_2_] decreased toward the middle of the mesocarp. As ripening progressed the minimum [O_2_] approached zero in the seeded cultivars and correlated to CD. Seed respiration was a large proportion of total berry respiration early in ripening but did not account for O_2_ deficiency late in ripening. [O_2_] increased towards the central axis corresponding to the presence of air spaces visualised using x-ray microCT. These connect to lenticels on the pedicel that were critical for berry O_2_ uptake as a function of temperature, and when blocked caused anoxia in the berry, ethanol accumulation and CD. Lenticel area on Chardonnay pedicels was higher than that for Shiraz probably accounting for the lower sensitivity of Chardonnay berry CD to high temperatures. The implications of hypoxia in grape berries are discussed in terms of its role in ripening and berry water relations.

**Highlight:** Grape berry internal oxygen concentration is dependent upon lenticels on the pedicel and cultivar differences in lenticels may account for temperature sensitivity of cell death in the mesocarp.

## Introduction

The onset and rate of cell death (CD) in the berry mesocarp of *Vitis vinifera* L are genotype-dependent, and modulated by temperature and drought (Bonada *et al.*, 2013a; Bonada *et al.*, 2013b; Fuentes *et al.*, 2010; Krasnow *et al.*, 2008; Tilbrook and Tyerman, 2008). Evolutionarily, cell death may have been selected as a trait that favours seed dispersal (Hardie *et al.*, 1996). Cell death correlates with berry dehydration (Bonada *et al.*, 2013b; Fuentes *et al.*, 2010), a common phenomenon in warm wine growing regions in Australia, and is partially distinct from other forms of “berry shrivel” (Bondada and Keller, 2012; Keller *et al.*, 2016). Berry dehydration associated with CD is common in Shiraz (Syrah), resulting in significant increases in sugar concentration (Caravia *et al.*, 2016; Rogiers *et al.*, 2004; Sadras and McCarthy, 2007). It is also associated with altered composition of fatty acids and anthocyanins, higher alcohol acetates, (Šuklje *et al.*, 2016), and alteration of the sensory characteristics of the berries at harvest (Bonada *et al.*, 2013a).

Grape berries are a non-climacteric fruit that do not exhibit a large rise in respiration rate or ethylene at the onset of ripening, though ethylene may still play a role (Bottcher *et al.*, 2013). However, the onset of ripening is associated with the accumulation of hydrogen peroxide (H_2_O_2_) in the skin of Pinot Noir berries, and there is increased catalase activity and peroxidation of galactolipids in skins (Pilati *et al.*, 2014). Although Pilati et al. (2014) considered that H_2_O_2_ was by a harmless signal, Pinot Noir also shows up to 50% CD later in ripening (Fuentes *et al.*, 2010). The accumulation of H_2_O_2_, apart from a potential signal for ripening (Pilati *et al.*, 2014), is also characteristic of plant tissues exposed to hypoxia or anoxia (Blokhina *et al.*, 2001; Fukao and Bailey-Serres, 2004). The respiratory quotient of grape berries increases during ripening (Harris *et al.*, 1971), in association with increased ethanolic fermentation (Famiani *et al.*, 2014; Terrier and Romieu, 2001) and suggests a major change in berry metabolism. Other fruit also show restricted aerobic respiration. For example in apple, tomato and chicory fruit, a clear effect of oxygen concentration [O_2_] on respiration and on the occurrence of fermentation was found (Hertog *et al.*, 1998). Ethanolic fermentation contributes to maintain cell function under O_2_-limiting conditions provided sugars are available. Interestingly, both H_2_O_2_ and ethylene have been implicated in its regulation (Fukao and Bailey-Serres, 2004).

Hypoxia-induced oxidative stress decreases lipid and membrane integrity (Blokhina *et al.*, 2001), the latter being clearly evident in most wine grape berries as indicated by vitality stains that depend on membrane integrity (Tilbrook and Tyerman, 2008). The decrease in cell vitality in Shiraz grapes is reflected by a decrease in extracellular electrical resistance during ripening, indicating leakage of electrolytes from cells (Caravia *et al.*, 2015). This leakage corresponds to the accumulation of potassium in the extracellular space of berries of Merlot (Keller and Shrestha, 2014) a cultivar that also undergoes CD (Fuentes *et al.*, 2010). O_2_ deprivation diminishes intracellular energy status that ultimately will challenge cell vitality in non-photosynthetic organs, as exemplified by roots under flooding or waterlogging (Voesenek *et al.*, 2006). Although the grape berry does show some photosynthesis in early stages of development (Ollat and Gaudillère, 1997), during ripening photosynthetic pigments and nitrogen content are reduced and atmospheric CO_2_ is not fixed while re-fixation of respiratory CO_2_ declines (Palliotti and Cartechini, 2001).

Shiraz berry CD can be accelerated by both water stress and elevated temperature (Bonada *et al.*, 2013a). There are increasing frequencies and intensities of heat waves and drought events globally and in Australia (Alexander and Arblaster, 2009; Perkins *et al.*, 2012) and the warming trend is predicted to have adverse effects on grapevine physiology (Webb *et al.*, 2007) and berry quality in warm regions (Bonada and Sadras, 2015; Caravia *et al.*, 2016; Fuentes *et al.*, 2010). CD is linked to high temperature stress in other plants. For example, when mustard seedlings were exposed over 1.5 h to 55 °C there was more than 2 folds increase in ROS production accompanied by the reduction in catalase activity, and cell death was triggered (Dat *et al.*, 1998). Higher temperature would also increase the demand for O_2_ to support increased oxidative respiration in the berry (Kriedemann, 1968). Meanwhile, O_2_ diffusion into the berry may be limited due to decreased gas exchange across the berry skin during ripening, as judged by declining transpiration (Rogiers *et al.*, 2004; Scharwies and Tyerman, 2017) and/or changes in berry internal porosity during ripening. Lenticels on the skin of potato tubers are the main channel for O_2_ uptake for respiration (Wigginton, 1973). The phellem-lenticel complex of woody roots and trunks also regulates transpiration and gas exchange including O_2_ (Lendzian, 2006). In the grape berry, the small number of stomata on the skin develop into non-functional lenticels occluded with wax during development (Rogiers *et al.*, 2004), but lenticels are very prominent on the pedicel (Becker *et al.*, 2012).

Wine-grape cultivars are seeded, and have been selected for wine-related attributes including pigments, and precursors of flavour and aroma whereas table-grape cultivars have been selected for turgor maintenance, and markets increasingly demand seedless fruit; these differences in selective pressures between wine and table grape cultivars have led to differences in the dynamics of water during berry ripening (Sadras *et al.*, 2008). Table grape seedless cultivars show little or no CD well into ripening (Fuentes *et al.*, 2010; Tilbrook and Tyerman, 2008). Although lignification of seeds is complete before berries begin to ripen (Cadot *et al.*, 2006), the oxidation of seed tannins is sustained (Ristic and Iland, 2005) and is concurrent with the oxidation of phenolic compounds such as flavan-3-ol monomers and procyanidins (Cadot *et al.*, 2006). Lignin polymerisation requires the consumption of O_2_ and generation of H_2_O_2_ for the final peroxidase reaction (Lee *et al.*, 2013), and this with oxidation of tannins could add additional stress to the mesocarp in seeded cultivars. Phenolic compounds can also act as ROS-scavengers (Blokhina *et al.*, 2003). In grape particularly, the biosynthesis of procyanidins coincide with the initial rapid period of berry growth (Coombe, 1973). Flavan-3-ol was shown to accumulate significantly during the early ripening stage (Cadot *et al.*, 2006). Taken together, seed respiration and maturation deserves consideration in understanding mesocarp cell death.

In this study we test the hypothesis that hypoxia occurs within the grape berry during ripening and that this may be correlated with the onset of cell death in the pericarp of seeded cultivars. We compared the patterns of CD and [O_2_] profiles across the berry flesh of two wine, seeded cultivars, Chardonnay and Shiraz, and a seedless table grape cultivar, Ruby Seedless. The respiratory demand of seeds and the berry were measured for different ripening stages and different temperatures. The diffusion pathway of O_2_ supply was assessed through examination of the role of lenticels on the pedicel of the berry and air space estimates using X-ray micro-computed tomography of single berries.

## Material and Methods

### Berries from vineyards

Details of sources of berries, sampling times and measurements are listed in Supplementary Table 1. Mature Shiraz, Chardonnay and Ruby Seedless vines on own roots were grown under standard vineyard management and irrigation at the Waite Campus (34°58'04.8"S 138°38'07.9"E), University of Adelaide Shiraz and Chardonnay, three replicates each consisted of 2 vines per replicate for Shiraz and 3 vines per replicate for Chardonnay. Ten random bunches were labelled within each replicate and 20 berries per replicate were carefully excised at the pedicel-rachis junction with sharp scissors at each sampling date. Ruby Seedless grapes were sampled from three vines with 5 bunches labelled for sampling on each vine and 20 berries were sampled from each vine. Timing of sampling during berry development was measured as days after anthesis (DAA, 50% of caps fallen from flowers). Berries were placed in sealed plastic bags into a cooled container, and taken to the laboratory where they were stored at 4 °C in the dark and tested within 48 hours of sampling. Berries harvested from the Waite vineyards were sampled over the 2014-2015, 2015-2016 and 2016-2017 seasons and were used for berry [O_2_] profile measurements and respiration measurements.

For visualizing berry internal air space, Shiraz berries from a vineyard in Nuriootpa, South Australia (34°28'32.9"S 139°00'28.0"E) were sampled during season 2015-2016 for x-ray micro-computed tomography where three berries, each from a different vine, were used for each sampling time

### Berries from pot-grown vines

Shiraz and Chardonnay cuttings were taken from the Waite vineyards in April 2015 and propagated after storage at 4 °C in the dark for approximately two weeks. Propagation method and vine nutrition management were based on Baby et al. (2014). Briefly, after roots were initiated in a heated sand bed in a 4 °C cold room for 8 weeks, and after the root length reached approximately 6 cm, cuttings were transferred into vermiculite: perlite (1:1) mixture in 12 cm pots. Pots were placed in a growth chamber with a 16 h photo-period, 400 μmol photons/(m^2^·s) at the plant level, 27 °C day/ 22°C night, and 50% humidity. Pots were irrigated with half strength Hoagland solution (Baby *et al.*, 2014). Fruitful vines at stage EL-12 (Coombe, 1995) were then transferred into a University of California (UC) soil mix: 61.5 L sand, 38.5 L peat moss, 50 g calcium hydroxide, 90 g calcium carbonate and 100 g Nitrophoska^®^ (12:5:1, N : P : K plus trace elements; Incitec Pivot Fertilisers, Southbank, Vic., Australia), per 100 L at pH 6.8, in 20 cm diameter (4 L) pots irrigated with water thereafter. Five berries (each from 3 different vines) of each cultivar were used for light stereomicroscopy.

Chardonnay rootlings were obtained from Yalumba Nursery in April 2017 and planted with UC mix soil and in the same growth chamber with the same growth conditions as above. Seven vines, each with one bunch, were used for O_2_ diffusion experiments.

### [O_2_] profiles in berries

Berry [O_2_] was measured using a Clark-type O_2_ microelectrode with a tip diameter of 25 µm (OX-25; Unisense A/S, Aarhus, Denmark). The microelectrodes were calibrated in a zero O_2_ solution (0.1M NaOH, 0.1M C6H7NaO6) and an aerated Milli-Q water (272 µmol/L at 22 °C), as 100% O_2_ solution. Individual berries (equilibrated to room temperature) were secured on the motorized micromanipulator stage. To aid the penetration of the microelectrode into the berry skin, the skin was pierced gently with a stainless-steel syringe needle (19G), to a depth of 0.2 mm, at the equator of the berry. The microsensor was positioned in the berry through this opening and [O_2_] profiles were taken with depth towards the centre of the berry. For Shiraz, measurements were taken from 0.2 mm to 1.5 mm under the skin at 0.1 mm increments. The electrode was not moved beyond this point to avoid damaging the tip against a seed. For Ruby Seedless and Chardonnay grapes, where there were no seeds present or the position of the seeds could be determined through the semi-transparent skin, measurements were taken at 0.5 mm intervals from 0.2 mm under the skin to the berry centre. Each measurement was applied for a 10s duration at each depth. Between each position, a 20s waiting time was applied to ensure stable signals. To test whether puncturing of the skin by the needle and insertion of the microelectrode contaminated the berry internal O_2_ by the surrounding air, a collar (plastic ring) was placed around the insertion site and a gentle stream (250 mL/min) of nitrogen gas was applied to the insertion point while obtaining the O_2_ readings (Fig 1A).These readings were compared to those where no nitrogen gas was applied.

**Fig. 1.**
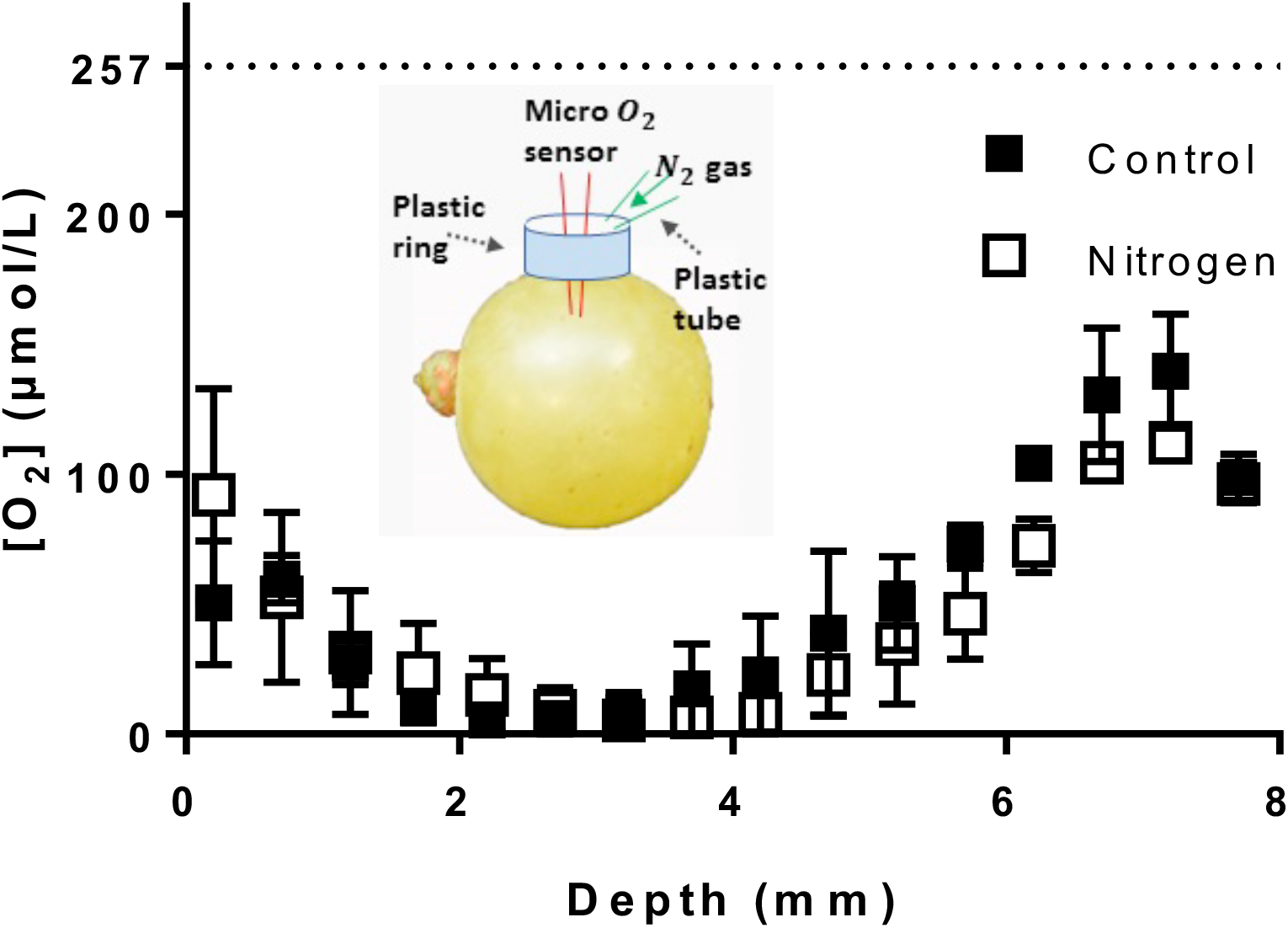
[O_2_] profiles of Chardonnay berries (90 DAA in season 2016-2017, Waite vineyards) measured with and without N_2_ gas applied at the entry point during measurement. Inset: experimental set-up for measuring berry [O_2_] profiles. Illustration not to scale. The O_2_ sensor (tip diameter 25 µm) was inserted at the equator of the berry and moved inwards to the centre approximately across the radius. Around the entry of the sensor, a plastic ring was sealed and glued to the berry, to contain nitrogen gas gently flowing on to the entry point of the sensor. Error bars are SEM (n=3). Two-way ANOVA (repeated) test showed depth accounted for 68.73% of total variation (p < 0.0001), treatments accounted for 0.55% of total variation (P = 0.26) and interaction accounted for 3.72% of total variation (P = 0.87).

The O_2_ readings were recorded using the Unisense Suite software (Unisense A/S, Aarhus, Denmark). Three berries were measured for each biological replicate. Means and SE of each step (n = 3) were calculated and [O_2_] profiles were compiled using GraphPad Prism 7 (GraphPad Software Inc., La Jolla, CA, USA). Following the O_2_ measurements, berry temperature was recorded using an IR thermometer (Fluke 568, Fluke Australia Pty Ltd, NSW, Australia) with a type-K thermocouple bead probe (Fluke 80PK-1). Berry diameters at the equator were measured with a digital calliper. Berry vitality was determined (see below) and total soluble solids (TSS) of the juice from individual berries was determined using a digital pocket refractometer (Atago, Tokyo, Japan) as an indicator of berry maturity.

### Testing the role of pedicel lenticels

[O_2_] were measured as above but with the probe stationary at approximately 2 mm from the pedicel along the berry central axis. After a stable reading was obtained N_2_ gas (250 mL/min) was then applied over the pedicel in order to test the contribution of pedicel lenticels to O_2_ diffusion into the berry.

### Berry and seed respiratory O_2_ consumption

A Clark-type oxygen microsensor OX-MR and the MicroRespiration System (Unisense A/S, Aarhus, Denmark) were used for berry and seed respiration measurements. A replicate consisted of 9 berries. The measuring chamber was filled with aerated MilliQ water, constantly stirred and was maintained at 25°C in a water bath. After the measurement of whole berry respiration, seeds of the 9 berries were extracted and the seed respiration rate measured using the same apparatus. Changes in the chamber’s water [O_2_] were monitored for at least 15 mins, with readings taken every 5 seconds in order to determine a steady respiration rate from the slope of the decline in [O_2_].

Respiration was also measured for Shiraz and Chardonnay berries before and after the pedicels were covered with silicone grease, at both 20 °C and 40 °C. Another batch of 9 Chardonnay berries was used to determine the respiratory contribution of excised pedicels.

The temperature dependence of berry respiration was determined with a water bath held at 10 °C, 20 °C, 30 °C and 40 °C.

### Pedicel lenticel density

The lenticel density of Chardonnay and Shiraz berry pedicels (stem and receptacle) was assessed using a Nikon SMZ 25 stereo microscope with CCD camera (Nikon Instruments Inc., Melville, NY, USA). Lenticel area (%) was estimated using ImageJ (Schneider *et al.*, 2012) by first adjusting the colour threshold of the image to separate the pedicel from the background and then the lenticels from the pedicel. Subsequently the ROI managing tool was used to estimate the relative area of the pedicels and the lenticels.

### Long term effect of blocking pedicel lenticels

The pedicel of approximately half of the berries on each bunch of growth chamber grown Chardonnay were covered with silicone grease at the onset of ripening (first signs of berry softening). Two or three pairs of berries, each pair containing one covered and one uncovered pedicel from one plant, were randomly sampled throughout the course of the experiment at 3, 5, 7, 10, 12, 14 and 18 days after application. Profiles of berry O_2_ concentration were measured as above, and berries were subsequently assessed for cell vitality (see below). Three pairs of berries were sampled 12 and 20 days after silicone application and assessed for internal ethanol concentration (see below).

### Berry ethanol concentration

A subsample of ten frozen berries were ground to a fine powder in a liquid nitrogen-cooled A11 basic mill (IKA, Germany). Ethanol was quantified using an Ethanol Assay kit following the manufacturer’s instructions (Megazyme International Ireland Ltd., Wicklow, Ireland). Briefly, alcohol dehydrogenase (ADH) catalysed the oxidation of ethanol to acetaldehyde. Acetaldehyde was then further oxidized to acetic acid and NADH in the presence of aldehyde dehydrogenase (AL-DH) and NAD^+^. NADH formation was measured in a FLUOstar Omega plate reader (BMG LABTECH GmbH, Ortenbery, Germany) at 340 nm.

### Pericarp cell vitality estimation

Cell vitality was estimated using a fluorescein diacetate (FDA) staining procedure on the cut medial longitudinal surface of berries (Fuentes *et al.*, 2010; Tilbrook and Tyerman, 2008). One half-berry was used to measure osmolality, while the cut surface of the other half was incubated in the dark for 15 min in a 4.8 µM FDA solution with the solution osmolality similar (to within 10%) of the grape juice (adjusted with sucrose). The stained berries were viewed under a Nikon SMZ 800 (Nikon Co., Toyko, Japan) dissecting microscope under ultraviolet light with a green fluorescent protein filter in place. Images were taken by a Nikon DS-5Mc digital camera (Tochigi Nikon Precision Co., Ltd, Otawara, Japan) and NIS-Elements F2.30 software with the same gain and exposure settings for all images. Images were analyzed with a MATLAB (Mathworks Inc., Natick, MA, USA) code for determining berry cell vitality (Fuentes *et al.*, 2010).

### Air spaces within the berry

Shiraz grapes were imaged at the x-ray micro computed tomography facility at Adelaide Microscopy, The University of Adelaide to map developmental changes in the distribution of air spaces within the berry tissues. Whole berries with pedicel attached were wrapped in foam sheets to secure the berries in the centre of the imaging chamber, and placed on the object tray of a Skyscan 1076 (Bruker microCT, Kontich, Belgium). 2D images were acquired with 59 kV source voltage, 149 uA source current, Al 0.5mm filter, 2356 ms exposure, 0.4-degree rotation step and 8.5 µm image pixel size. The 2D projection images were reconstructed into stacks of transverse images using NRecon (bruker-microct.com). Berry air porosity was then estimated with CTan software (bruker-microct.com) with a custom plugin using the transverse images. 3D images of the berry internal air space were generated using CTvox software (bruker-microct.com), using colour rendering modules to distinguish the air porous volume from the berry volume. These two different colour rendering schemes were then aligned together to create images showing air pore distribution inside the berries.

### Statistical analysis

All data are presented as mean ± SEM. The effect of O_2_ sensor depth and applying N_2_ gas at the point of sensor entry on [O_2_], the effect of O_2_ sensor depth and ripening stage of berries on [O_2_], the effect of temperature and covering lenticels on respiration, the effect of temperature and grape maturity on respiratory Q_10_, the effect of covering lenticels and the duration of coverage on [O_2_] and the effect of covering lenticels and ripening stage on ethanol in Chardonnay berries were all tested with two-way ANOVA. Deming regression was used to determine the association between fluorescent intensity (grey value) and [O_2_], this type of regression takes account for error in both x and y (Strike, 1991). Significance of differences in respiration of berry and seed of Chardonnay at two ripening stages, significance of differences of lenticel area on pedicels between Chardonnay and Shiraz, significance of differences of activation energy of O_2_ uptake of Chardonnay and Shiraz berries each at two ripening stages and significance of differences of porosity and connectivity index in Shiraz at two ripening stages were all tested with t-tests. Second order polynomial regression was used to determine the association between TSS/sugar per berry and days after covering lenticels. Cell death rates in lenticel covered berries and control berries were determined using linear regression.

## Results

### Internal oxygen profiles of grape berries

In Chardonnay, [O_2_] decreased from the skin towards the interior of the mesocarp to reach very low concentration at depths of 2.2 mm to 4 mm (Fig. 1). The minimum [O_2_] over this depth range was 5.5 ± 5.5 µmol/L and thus could be considered bordering on anoxic. However, with further penetration towards the central axis of the berry, [O_2_] increased and reached a maximum at a depth of 7 mm (Fig. 1). To test if the [O_2_] profiles were affected by introduced O_2_ via the penetration site through the skin, N_2_ gas was gently applied on to the entry point of the sensor during the measurements. The [O_2_] profiles were similar for control and nitrogen-treated berries (Fig. 1) indicating that we could exclude leakage through the site of penetration as a significant factor determining the recorded profiles.

### Progression of cell death and changes in internal oxygen profiles of grape berries

To determine if there was a link between the progression of cell death and hypoxia within the berry we determined CD using the FDA vital staining technique and recorded [O_2_] profiles on the same batches of berries at different development stages. Similar [O_2_] profiles were observed for Chardonnay and Ruby Seedless (Fig. 2A, C), and for Shiraz over the first 1.5 mm (Fig. 2E), but the [O_2_] dropped more steeply across the skins as ripening progressed in all cultivars resulting in overall lower [O_2_] across the berry. This was manifest as much lower minimum [O_2_] at the last ripening stage sampled: Chardonnay 0 µmol/L, Ruby Seedless 14.9 ± 8.86 µmol/L, Shiraz 0 µmol/L. Because seeds could not be visualised in Shiraz berries the micro oxygen sensor could not be moved further into the berry than about 1.6 mm without risking the integrity of the sensor (Fig. 2E). Nevertheless, it was clear that [O_2_] dropped precipitously towards 1 mm (Fig. 2E).

**Fig. 2.**
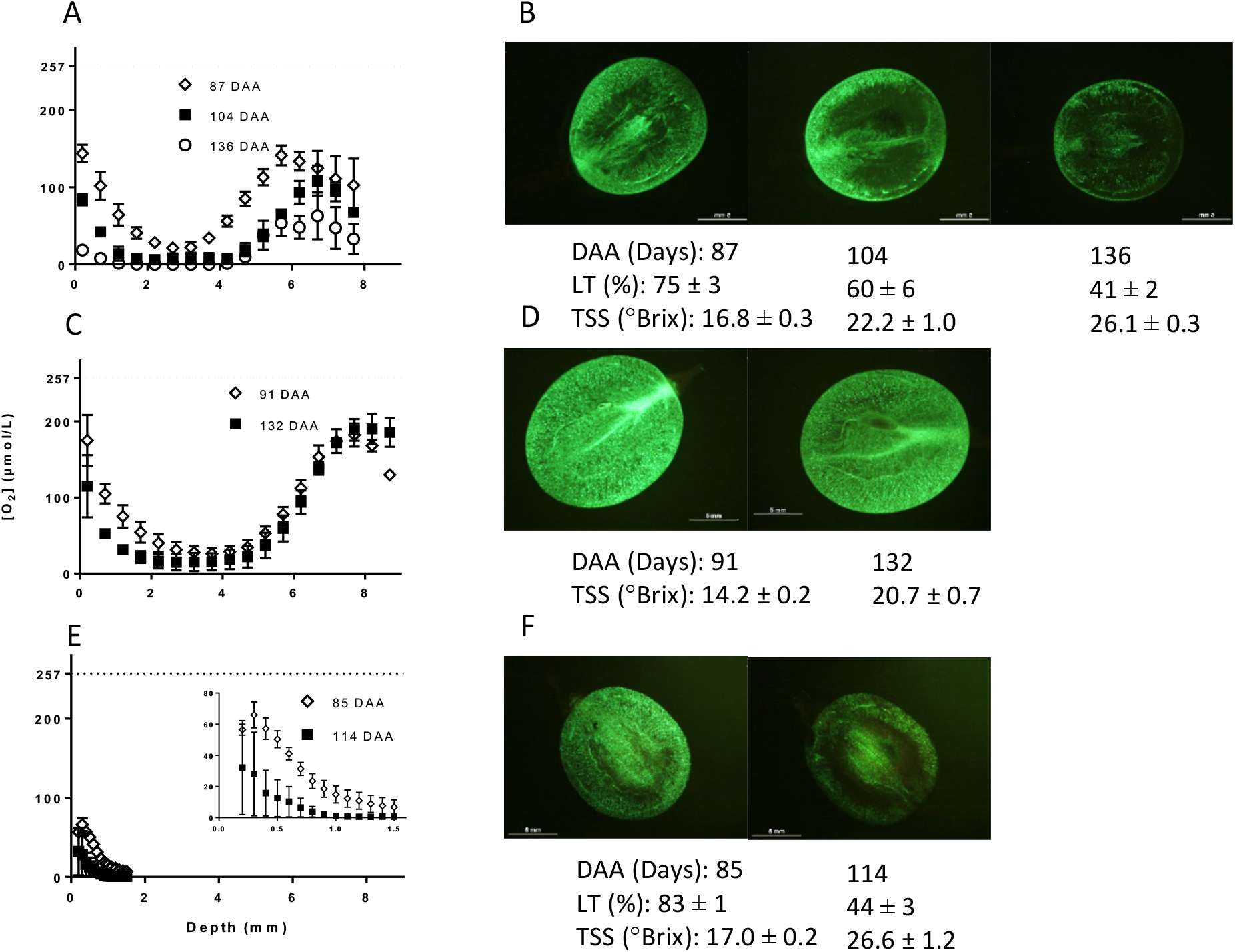
[O_2_] profiles of Chardonnay, Ruby Seedless and Shiraz berries at various ripening stages and examples of living tissue (LT) in the pericarp (Waite vineyards). (A) Chardonnay berries were sampled at 87, 104 and 136 DAA in 2015-2016 season. Two-way ANOVA (repeated) test showed depth accounted for 46.75% of total variation (p < 0.0001), time accounted for 29.93% of total variation (P < 0.0001) and interaction accounted for 8.029% of total variation (P = 0.058). Horizontal dashed line indicates the approximate O_2_ saturation value for Millipore water at room temperature, same as berries at the time of measurement. (B) Images of medial longitudinal sections stained with FDA hi-lighting LT differences at different stages of ripening. (C) [O_2_] profiles of Ruby Seedless berries sampled at 91 and 132 DAA in 2016-2017 season. Two-way ANOVA (repeated) test showed depth accounted for 85.22% of total variation (p < 0.0001), time accounted for 1.2% of total variation (P = 0.0025) and interaction accounted for 3.731% of total variation (P = 0.048). (D) LT was close to 100% for the two respective sampling days. Two-way ANOVA (repeated) test showed depth accounted for 40.86% of total variation (p = 0.0005), time accounted for 19.57% of total variation (P < 0.0001) and interaction accounted for 6.39% of total variation (P = 0.43). (E) [O_2_] profiles of Shiraz berries sampled on 85 and 114 DAA in 2014-2015 season. Error bars are SEM (n = 3) for A, C and E.

Vitality staining (Fig. 2B, F) indicated that, for both Chardonnay and Shiraz, living tissue decreased over time as TSS accumulated and occurred predominately in the middle of the mesocarp corresponding to the minimum in [O_2_]. Further, the change in fluorescent signal intensity across the radius at the equator of Chardonnay berries showed a similar trend as for berry internal [O_2_] (Fig. 3A), indicating a correlation between cell vitality and internal [O_2_] (Fig. 3B). On the other hand, Ruby Seedless berries maintained cell vitality close to 100% up to 132 DAA, when TSS was 20.7 °Brix (Fig. 2D). While a similar shape of [O_2_] profile was observed within the mesocarp of Ruby Seedless berries when compared with that of Chardonnay berries (Fig. 2C), [O_2_] did not reach zero.

**Fig. 3.**
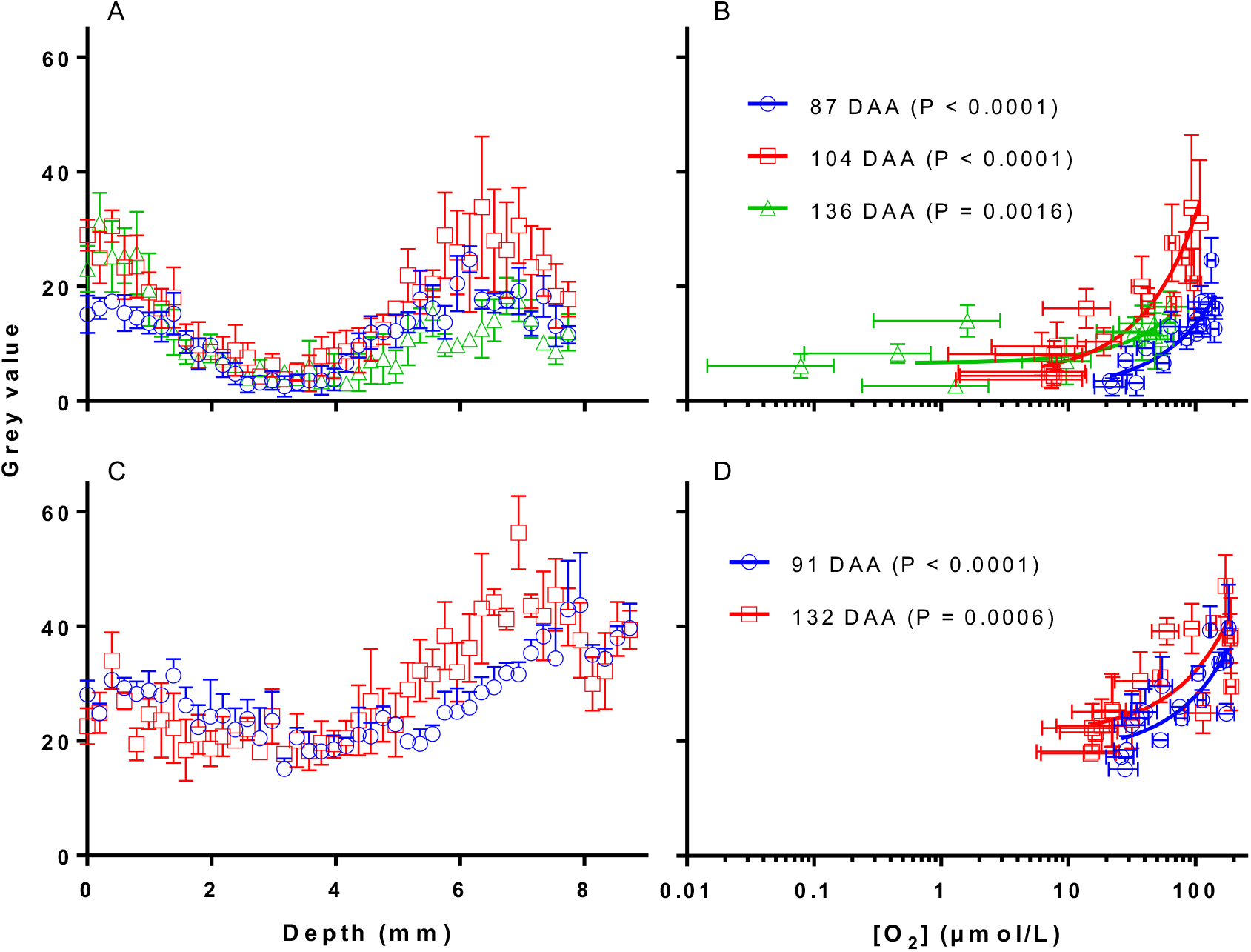
Correlation between berry living tissue and [O_2_]. Fluorescent signal intensity (grey value) across radius at equator of Chardonnay (A) and Ruby Seedless (C). Correlation (Deming regression) between fluorescent signal intensity and [O_2_] at corresponding depths in Chardonnay (B) and Ruby Seedless (D) ([O_2_] profiles shown in Fig. 1). Error bars are SEM (n=3).

Despite the decrease in oxygen concentration across the mesocarp over ripening, for Chardonnay and Ruby Seedless berries, [O_2_] started to increase with depth from about 4.2 mm and reached a maximum at around 6.2 mm in Chardonnay and 8.2 mm in the larger Ruby Seedless berries (Fig. 2A, C). Standardising the position of the sensor relative to the diameter of each berry replicate (Fig. 4), showed that [O_2_] peaked at the central vascular bundle region at all sampling times for both Chardonnay (Fig. 4A) and Ruby Seedless (Fig. 4B).

**Fig. 4.**
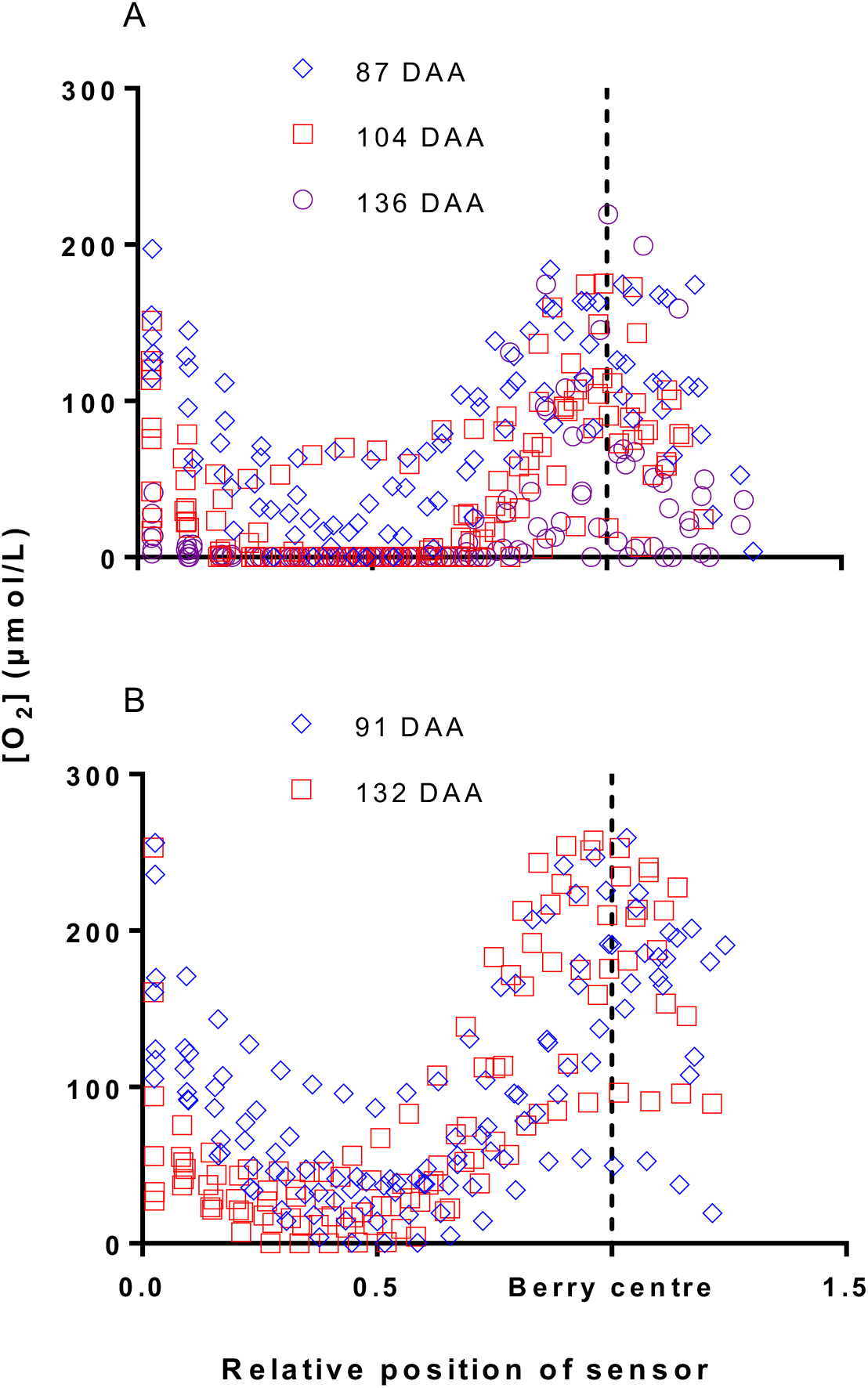
Individual berry [O_2_] profiles normalized to the berry radii that differed between replicates. (A) [O_2_] profiles of Chardonnay berries sampled at 87, 104 and 136 DAA in 2015-2016 season (Mean data shown in Fig. 2). (B) [O_2_] profiles of individual Ruby Seedless berries sampled at 91 and 132 DAA in 2016-2017 season.

### Consumption and supply pathways of oxygen within grape berries

Considering the link between living tissue (%) and [O_2_] (Fig. 3), and the maintenance of living tissue (%) in well-developed berries of Ruby Seedless (Fig. 2D), we investigated the contribution of seeds to the respiratory demand of the berry in Chardonnay. Seed fresh weight peaks at the beginning of sugar accumulation and skin coloration (termed veraison) (Ristic and Iland, 2005) (around 63 DAA for Chardonnay here). Seed respiration at this stage was 5-fold higher than whole berry respiration on a per gram basis. Berry respiration on a per gram basis reduced by about a third at 122 DAA compared to 63 DAA (Fig. 5A), however seed respiration decreased by 40-fold (Fig. 5B). Berry mass nearly doubled from 7.2 ± 0.5 g at 63 DAA to 13.9 ± 1.4 g 122 DAA, thus on a per berry basis respiration rate increased by about 18% from 63 DAA to 122 DAA (Fig. 5C). The contribution from the total number of seeds in the berry accounted for more than half of the respiratory demand in berries at veraison. This dropped to an insignificant proportion at 122 DAA (Fig. 5C).

**Fig. 5.**
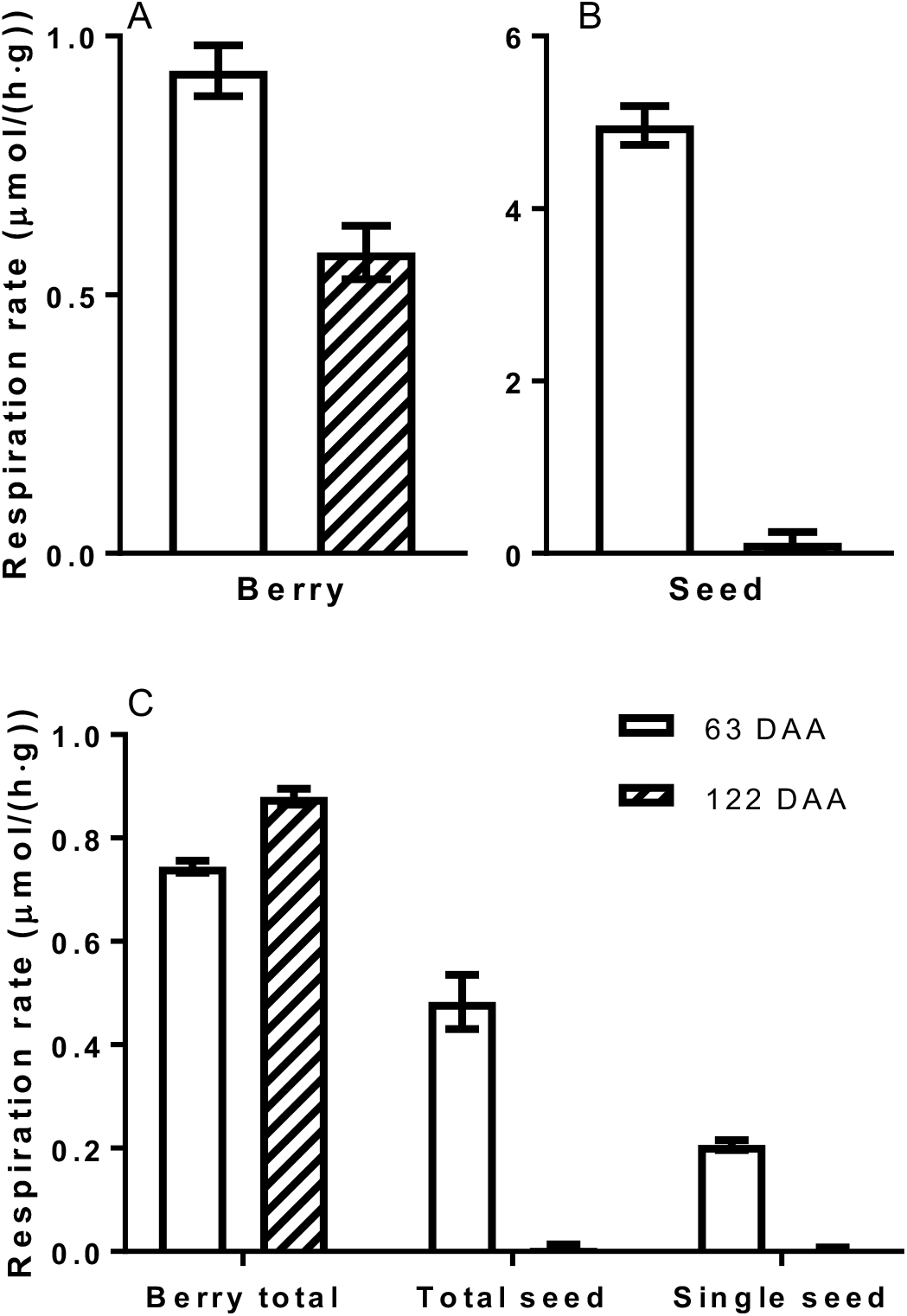
Chardonnay berry and seed respiration (25 °C) at 63 and 122 DAA in 2015-2016 season (Waite vineyards), illustrating the dramatic change in seed respiration. Respiration on a per gram basis for berries (A) and seeds (B). (C) Comparison of berry total (including seeds), all seeds per berry and single seed respiration. Error bars SEM (n = 3). All rates are different between 63 and 122 DAA (t-test, P < 0.05).

Differences in resistance to diffusion into the berry may influence the [O_2_] profiles. The pedicel lenticels may offer a pathway for O_2_ entry that could account for the higher concentration towards the central axis of the berry. There were obvious differences in lenticel morphology between Chardonnay (Fig. 6A) and Shiraz berries (Fig. 6B). Individual lenticels on Chardonnay pedicels were larger, and also had 10-fold larger total surface area as a proportion of pedicel surface area compared to that of Shiraz berries (Fig. 6C).

**Fig. 6.**
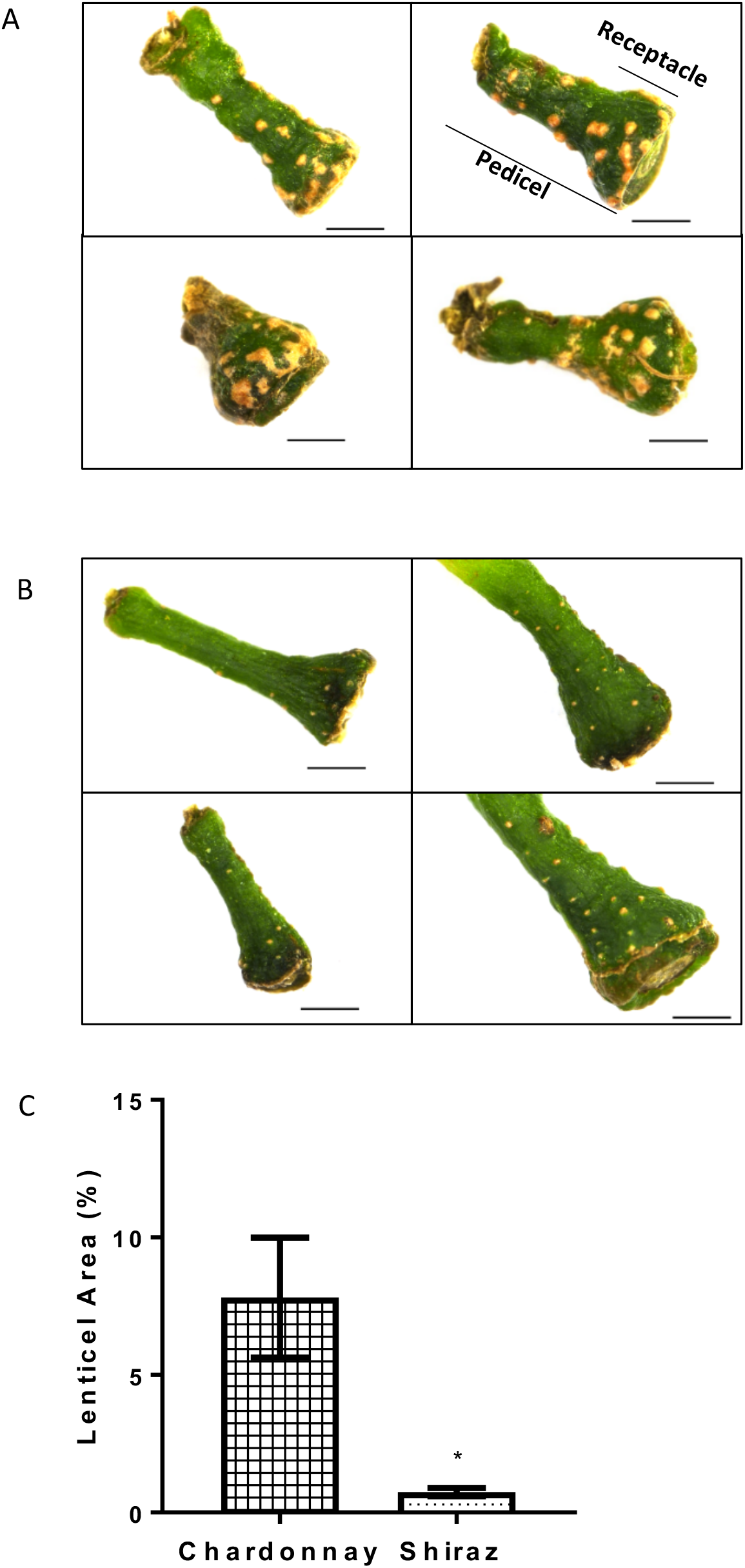
Differences in lenticel morphology and relative lenticel area between Chardonnay (A) and Shiraz (B) berry pedicels. (C) Lenticel area relative to pedicel surface area of Chardonnay and Shiraz berries (chamber grown, 2015) estimated using ImageJ. Error bars SEM (n = 5). *Significantly different (t-test, P < 0.05). Scale bars = 1mm.

To determine whether lenticels on the pedicel could be sites for berry gas exchange, respiration was measured on the same batches of berries with or without pedicels covered with silicone grease to impede gas exchange. This was examined at 20 and 40 °C as respiratory demand for O_2_ increases with temperature (Hertog *et al.*, 1998). Fig. 7A shows that covering the berry pedicel with the silicone decreased berry respiration rate at 40 ºC for both Shiraz and Chardonnay berries, but had no effect on respiration rate at 20 ºC. The temperature dependence of respiration was examined in more detail for Chardonnay and Shiraz with both yielding similar activation energies (Supplementary Fig. S1) that did not differ between berries sampled on the two days for each cultivar. The Q_10_ of Chardonnay (Supplementary Fig. S2) obtained at different temperature ranges (10 °C difference between 10 and 40 °C) showed no difference for berries sampled on the two days, nor were these values different between the two days over the same temperature range (Supplementary Fig. S2). The Q_10_ of Shiraz berries on 71 DAA (Supplementary Fig. S2) was the same across the temperature range. However, the Q_10_ of Shiraz berries on 113 DAA was higher for the 20-30 °C range relative to the other temperature ranges (Supplementary Fig. S2). The Q_10_ of Shiraz berries on 71 DAA also differed from the Q_10_ of berries on 113 DAA at the10-20 and 20-30 °C temperature classes. The decreased apparent respiration of berries with the coated pedicel was not due to the elimination of pedicel respiration because pedicel respiration rate at 40°C was a small fraction of the total berry respiration (Fig. 7B) and did not account for the decrease observed when pedicels were covered (Fig. 7A), where the decrease in respiration of pedicel-coated Shiraz and Chardonnay was 839.7 ± 101.8 and 1233.9 ± 229.4 nmol/h per berry, at 40 °C.

**Fig. 7.**
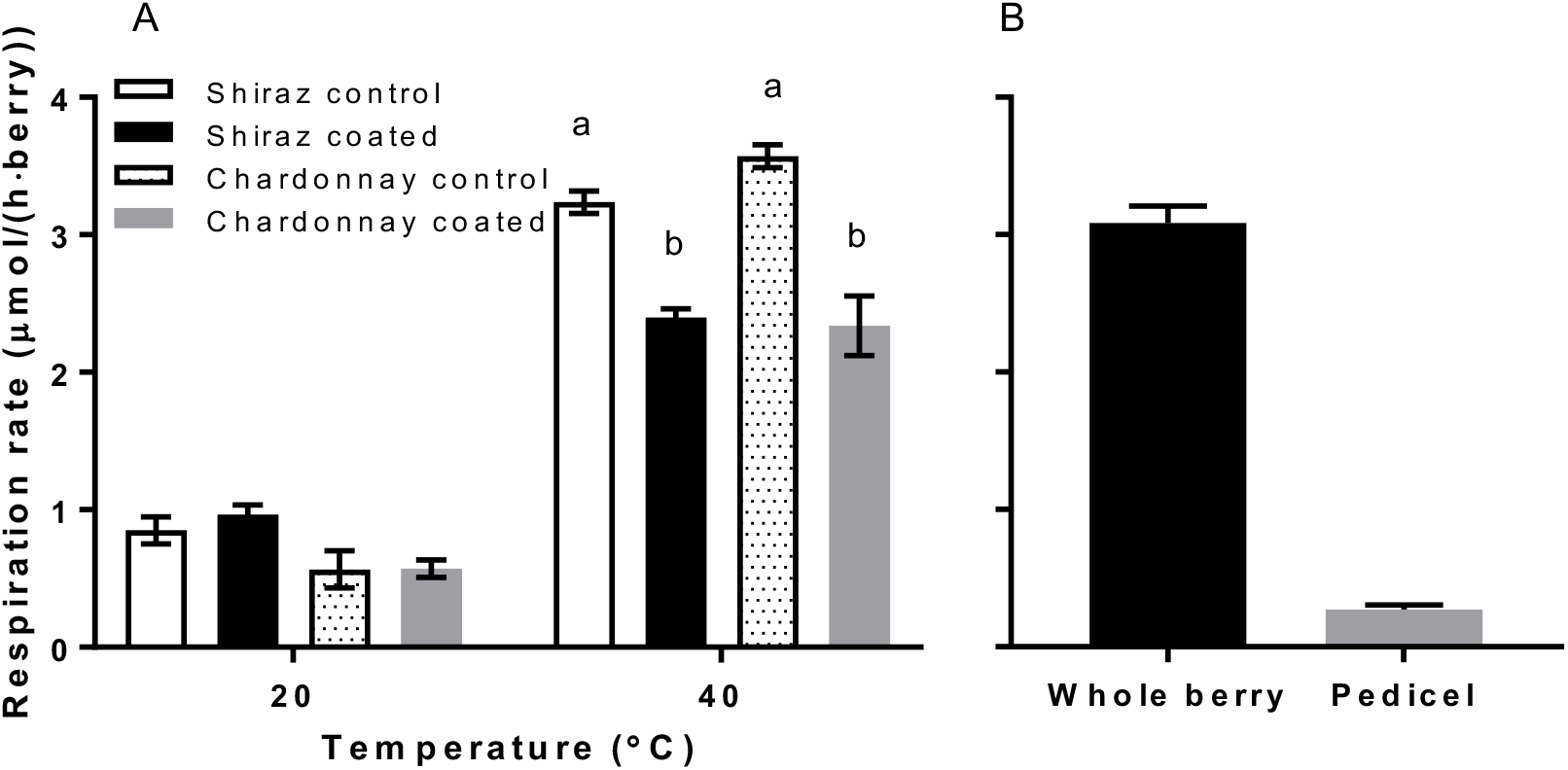
Role of the pedicel in oxygen diffusion as a function of temperature. (A) Respiration of Chardonnay (86 DAA) and Shiraz (77 DAA) berries at 20 and 40 °C with pedicels attached (2016-2017 season, Waite vineyards). Silicone grease was applied over the lenticels on the pedicel and receptacle regions (coated berries). At 20°C no significant difference in apparent berry respiration was found between control and pedicel coated berries for both cultivars. At 40°C both cultivars showed significantly lower apparent respiration compared to respiration of non-coated berries. Shiraz and Chardonnay each showed a decrease of 839.7 ± 101.8 and 1377.3 ± 161.3 nmol/hour per berry in respiration at 40°C (26 and 39% decrease) respectively. Different lower-case letters indicate significant difference between treatments at 40°C within each cultivar (two-way ANOVA, P < 0.0001). (B) Respiration rate of whole berry including attached pedicel and respiration of separated pedicels for Chardonnay at 40°C. The pedicel accounted for 9% of the whole berry and pedicel respiration rate. Error bars SEM (n = 3).

A rapid decrease in [O_2_] was observed at approximately 2 mm away from pedicel and close to the centre axis in the Ruby Seedless berries, when a N_2_ stream was activated over the pedicel (Fig. 8B).

**Fig. 8.**
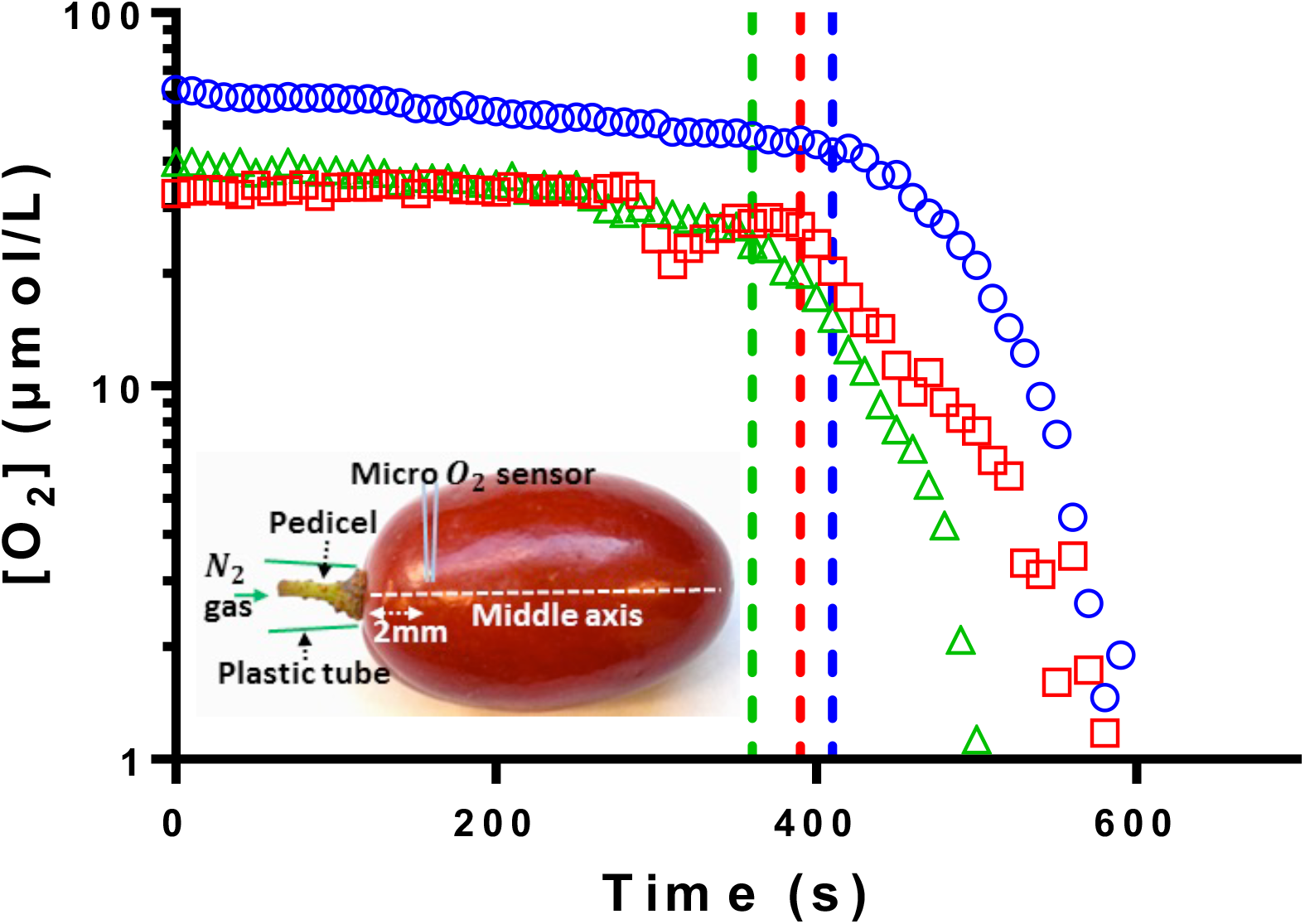
The role of the pedicel in gas diffusion into Ruby Seedless grapes (132 DAA in 2016-2017 season, Waite vineyards). [O_2_] of three individual berries as a function of time with the sensor inserted approximately at the central axis of Ruby seedless around 2 mm from pedicel. Dashed lines indicate the start of external N_2_ gas delivery over the pedicel. Inset: experimental set-up for applying N_2_ gas over the berry pedicel while measuring [O_2_].

An experiment was subsequently conducted using growth chamber grown Chardonnay vines to test whether blocking the lenticels of berries still attached to the vine would affect internal [O_2_] profiles. Three days after covering the berry pedicel with the silicone grease, a reduction in [O_2_] at the central vascular region was evident and remained at 0 µmol/L over the subsequent 15 days (Fig. 9A). For the uncovered (control) berries, a maximum of [O_2_] was evident at the central axis across all the days of measurement. Note that for each measurement day a different set of berries were assessed. Berry ethanol concentration of berries was measured at 12 and 20 days after blocking the lenticels. Lenticel blocked berries showed higher ethanol content compared to unblocked berries (Fig. 9B) consistent with a greater degree of fermentation within the hypoxic berries. Concentration of total soluble solids increased with time during the course of this experiment, and was higher for lenticel covered berries (Fig. 9C). Sugar/berry was not affected by covering the lenticel (Fig. 9D). Cell death was significantly increased by limiting oxygen diffusion after 10 days of covering the lenticels (Fig. 9E).

**Fig. 9.**
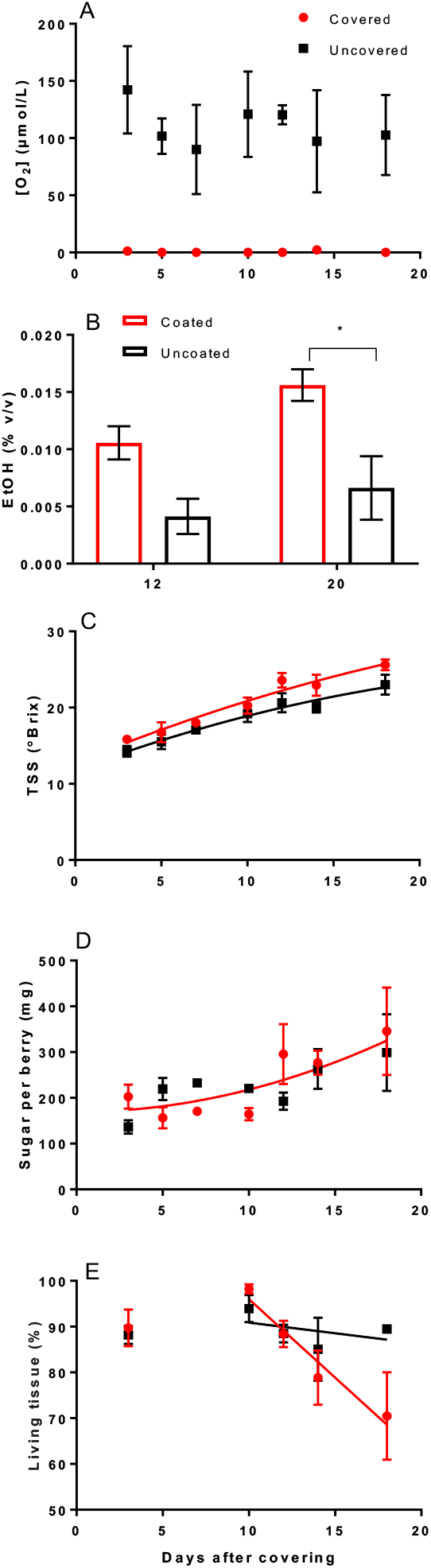
Blocking lenticels on berry pedicels changed O_2_ availability within Chardonnay berries (chamber grown, 2017). (A) [O_2_] at approximate centre axis of berries. Two-way ANOVA showed covering pedicels reduced [O_2_] (P < 0.0001). Time (P = 0.92) and interaction (P = 0.93) effects were not significant. (B) Ethanol content of berries after 12 and 20 days with (red)/without (black) silicone grease covering the berry pedicels. Two-way ANOVA (Tukey’s multiple comparisons test) showed coating the pedicel increased berry EtOH content at 20 days after coating (P = 0.036). (C) Pedicel covered berries showed significantly higher TSS during the course of the experiment compare to the uncovered berries (Second order polynomial, F-test, P = 0.0055). (D) No significant difference was found between treatments in sugar/berry (Second order polynomial, F-test, P = 0.64). (E) Significant decline of cell vitality found for covered berries. Slope of fitted line for covered berries is non-zero (P = 0.025) and different from slope of fitted line for uncovered berries. Error bars SEM. For B, n = 3. For A, C and D, n = 3 on 3, 5 and 18 days, and n = 2 on 7, 10 and 18 days.

### Air spaces within the grape berry shown by micro computed tomography (micro-CT)

Using micro-CT, the internal air spaces of Shiraz berries were visualized without any disruption at two time points during ripening. Application of a protocol that highlights airspace within the berries and colour codes the size of the airspace is shown in Fig. 10. There were air channels (delineated white) connecting the proximal region of the berry and the cavities around seeds in berries sampled on both 76 DAA (Fig. 10B) and 133 DAA (Fig. 10D) berries. White delineated regions indicated larger continuous air volume. In berries sampled on both days, it was evident that there were continuous air channels connecting the pedicel to the locule around the berry seeds. The dark blue colour highlighted the smaller air spaces in the mesocarp, which appeared to be denser in the berry sampled on 76 DAA than in the berry sampled on 133 DAA. Quantitative analysis was performed in the berry tissue region between the receptacle and the top (hilum) of the seeds. Both analysis of total porosity and connectivity of the air space, pores and channels, showed no significant differences between Shiraz berries sampled on the two days (Supplementary Fig. S3).

**Fig. 10.**
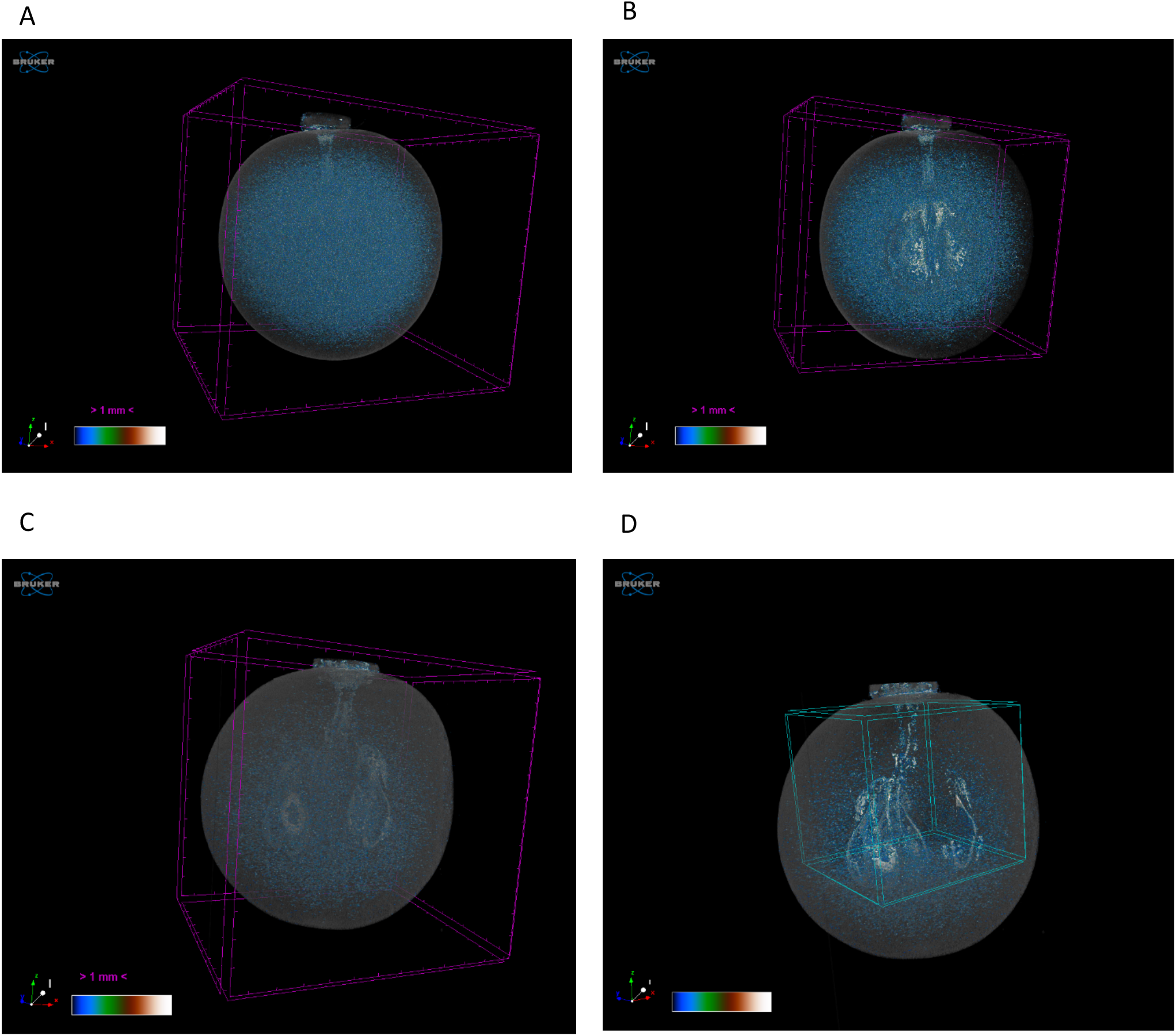
Micro CT 3D models of air spaces in Shiraz berries sampled on 76 DAA (A, B) (19.3 °Brix) and 133 DAA (C, D) (29.1 °Brix) in 2015-2016 season. Colour bar indicates magnitude of volumes highlighted, dark/blue to white colour represents smaller to larger volume. Visible volumes are within purple scaled frame (A, B, C). For (B), length of y-axis was shortened to reveal more details around the seeds. Blue cubic frames (D) removed the volume within, to reveal the structural arrangement of the air space.

## Discussion

The mesocarp of seeded wine grape berries typically shows a type of programmed cell death associated with dehydration and flavour development late in ripening (Fuentes *et al.*, 2010) (Bonada *et al.*, 2013a; Tilbrook and Tyerman, 2008). Here we show a close similarity between the pattern of CD across the berry mesocarp and [O_2_] profiles where the central regions of the mesocarp had both the highest CD and the lowest [O_2_]. In both Shiraz and Chardonnay the oxygen deficit in the centre of the mesocarp increased as ripening and cell death progressed, essentially becoming anoxic after about 100 days from anthesis under our experimental conditions. This contrasted to the seedless, table-grape cultivar where O_2_ concentrations remained above about 15 µmol L^−1^ (1.1 kPa) in the mid region of the mesocarp, still considered to be hypoxic (Saglio *et al.*, 1988), where CD was less apparent. In our experimental system, however, only three cultivars were tested and there is a confounded effect between cultivar types (wine vs table) with different water and sugar dynamics (Sadras *et al.*, 2008) and between seeded and seedless types. Separating these effects would require the comparison of seeded and seedless isogenic lines. Nonetheless, the strong correlation between CD and [O_2_] profiles, the role of lenticels, seed respiration, ethanol fermentation and CT-images all converge to support our working hypothesis that hypoxia or anoxia in the mesocarp contributes to CD in the grape berry.

The minimum [O_2_] we measured in the pericarp for both Chardonnay and Shiraz berries (close to zero) may be at or below the K_m_ for cytochrome C oxidase (0.14 μM) (Millar *et al.*, 1994), and very likely resulted in restricted oxidative phosphorylation and a shift to fermentation as evidenced by the detection of ethanol in Chardonnay berries; testing other cultivars for ethanol production would be of interest. All aerobic organisms require O_2_ for efficient ATP production through oxidative phosphorylation. Lower ATP production occurs under hypoxia when cells shift from oxidative phosphorylation to fermentation (Drew, 1997; Geigenberger, 2003; Ricard *et al.*, 1994). The depletion of ATP has profound consequences on cell physiology, including a change in energy consumption and cellular metabolism (Bailey-Serres and Chang, 2005; Drew, 1997). Loss of membrane integrity responsible for browning disorder in pears is also linked to internal hypoxia and low ATP levels (Franck *et al.*, 2007; Saquet *et al.*, 2003).

Survival of grape berry mitochondria after imposed anaerobiosis (based on succinate oxidation rates) is cultivar dependent with survival ranging from 1 to 10 days (Romieu *et al.*, 1992). This work was based on the process of carbonic maceration, a wine making procedure where whole berries ferment in an anaerobic atmosphere prior to crushing. Ethanol alters the respiratory quotient of grape mitochondria and uncouples oxidative phosphorylation (Romieu *et al.*, 1992). These effects occurred above 1% (vol) ethanol and well above the concentrations we measured in Chardonnay berries (0.015%); however it is possible for locally high concentrations of ethanol within the berry in our case. In a later paper, alcohol dehydrogenase (ADH) activity and ADH RNA were found to be already high in field grown Chardonnay berries before anaerobiosis treatment, which suggested a hypoxic situation already existed in the grapes as a result of some stressful conditions in the field (Tesnière *et al.*, 1993). Our results show that this may be the norm for certain regions within the berry mesocarp and likely exacerbated by high temperature (see below).

The internal [O_2_] of fruit depends on the respiratory demand, and the O_2_ diffusion properties of the skin and internal tissues. These can show genotypic differences as is the case for apple fruit (Ho *et al.*, 2010). In pear fruit differences in porosity of the cortex, the connectivity of intercellular spaces and cell distribution may account for variation between cultivars (Ho *et al.*, 2009). For pear it was possible to reconcile the observed variation in gas diffusion with the irregular microstructure of the tissue using a microscale model of gas diffusion. This also appears to be the case for different cultivars of apple as assessed by micro-CT (Mendoza *et al.*, 2007). For grape berries the [O_2_] profiles in our study would suggest a very low O_2_ diffusivity for the skin since a steep gradient occurred across the skin. Apple skin also showed a very low O_2_ diffusivity and likewise a steep concentration gradient across the skin (Ho *et al.*, 2010). Since sub-skin [O_2_] of grape berries declined dramatically during ripening for all three grape cultivars it would suggest a decline in O_2_ diffusivity during ripening that may result from the same epidermal and cuticle structural changes that cause a decline in berry transpiration (Rogiers *et al.*, 2004).

Changing properties of the skin, berry porosity and lenticels in the pedicel may all contribute to the reduced internal [O_2_] in grape berries during ripening. Fruit parenchyma can be regarded as a porous medium with air spaces distributed in between the elliptically tessellated cells (Gray *et al.*, 1999; Herremans *et al.*, 2015; Mebatsion *et al.*, 2006). We observed dense small air pores in the mesocarp of younger Shiraz berries that decreased in density in more mature berries corresponding with the decrease in [O_2_]. A maximum [O_2_] at the central axis region of both seeded and seedless berries throughout berry development, indicates a channel connecting the source of O_2_ intake and the central vascular bundles. Using different approaches, including blockage of pedicel lenticels with silicone grease or applying of N_2_ over pedicels, our experiments demonstrated that the pedicel lenticels are a major pathway for O_2_ diffusion into the grape berry. This corresponds to the predominant air canals observed in micro-CT from the receptacle into the central axis of the berry. Micro-CT to study air space distributions in fruit can reveal important properties that affect gas diffusion (Herremans *et al.*, 2015; Mendoza *et al.*, 2010) and can reveal internal disorders (Lammertyn *et al.*, 2003). In our work the visualisation of air space connecting the lenticels on the pedicel with the locular cavity around seeds provides the structural link to the measured peaks in [O_2_] around the central vascular region in the berries. This also confirmed the potential O_2_ uptake pathway through the pedicel lenticels, and distribution through the vascular networks. The relatively higher [O_2_] around both central and peripheral vascular bundles may be important for maintaining phloem unloading in the berry, and it is interesting to note that even with severe CD in berries the vascular bundles generally remain vital (Fuentes *et al.*, 2010). Despite this we observed higher sugar concentrations in essentially anoxic berries that had their lenticels blocked while still on the vine. This anomaly may be accounted for by decreased water influx as a result of anoxia causing an increase in sugar concentration.

Lenticels are multicellular structures produced from phellogen that replace stomata after secondary growth (Lendzian, 2006). The impact of lenticels on gas and water permeance compared to periderm of stems has been obtained for some species. For *Betula pendula*, the presence of lenticels substantially increased the water permeability of the periderm by between 26 and 53-fold (Schonherr and Ziegler, 1980). Lenticels on the berry pedicel are a preferential site for water uptake for submerged detached berries (Becker *et al.*, 2012). Water vapour and O_2_ permeance of tree phellem with and without lenticels showed that lenticels increased O_2_ permeance much more than that for water, over 1000-fold for one species, yet the permeance for water vapour was higher than that for O_2_ (Groh *et al.*, 2002). Interestingly, Schonherr and Ziegler (1980) showed that as the water vapour activity declined (increased vapour pressure deficit), water permeability was strongly reduced. If declining water vapour activity also reduced O_2_ permeability in grape berry lenticels this could restrictO_2_ diffusion under the very conditions where respiratory demand is increased, i.e. under water stress and with high temperature and vapour pressure deficit.

The decrease in [O_2_] at the approximate central axis in the seeded Chardonnay berry during development suggests there could be either an increase in respiratory demand, a decrease in the intake of O_2_ via the pedicel lenticels or decreased porosity through the central proximal axis. Ruby Seedless berries on the other hand did not show this reduction. This indicates there could be structural differences in lenticels between the seeded wine grape cultivar and the seedless table grape, or that the seeds themselves become a significant O_2_ sink (unlikely based on the arguments presented below). The lower lenticel surface area in Shiraz could be indicative of a greater restriction to O_2_ diffusion compared to Chardonnay. Shiraz is well known for its earlier and more rapid increase in CD under warm conditions (Bonada *et al.*, 2013b; Fuentes *et al.*, 2010). Unfortunately, it was not possible for us to probe for [O_2_] in the central region of the Shiraz berry to compare with Chardonnay berries. There appeared to be no detectable reduction in connectivity index or porosity of the proximal region of the berry between seeds and pedicel in Shiraz during ripening. The role of the pedicel lenticels in allowing grape berries to “breathe” and their variation between cultivars seems to have been overlooked and appears to be unique amongst fruit. Bunch compactness and pedicel length could also affect the gas diffusion via this passage, ultimately resulting in differences in berry internal oxygen availability throughout ripening.

Another possible explanation for the difference in oxygen profiles between the seeded and seedless cultivars is that seeds are a significant O_2_ sink late in ripening. Oxygen supply to seeds is essential for seed growth, and deposition of protein and oil (Borisjuk and Rolletschek, 2009). On the other hand, low [O_2_] within seeds favours low levels of ROS thus preventing cellular damage (Simontacchi *et al.*, 1995). The seeded win grape cultivars Riesling and Bastardo, increased O_2_ uptake from less than 0.45 µmol/h per berry to approximately 3 µmol/h per berry during early ripening, contrasting to seedless Sultana where the maximum O_2_ uptake was 1.5 µmol/h per berry (Harris *et al.*, 1971). We observed that total seed respiration was more than half of whole berry respiration at around the beginning of ripening. This high O_2_ demand from seeds, prior to the lignification of the outer layer (Cadot *et al.*, 2006), may create a significant O_2_ demand within the berry that could lower O_2_ concentrations in the locule, and potentially lowering the [O_2_] in the mesocarp. However, seed respiration in Chardonnay dramatically declined later in ripening, accounting for the decrease in berry respiration on a per gram basis. During late ripening, [O_2_] in the mesocarp of the seeded cultivar dropped to almost zero. Therefore, it is unlikely that the lower [O_2_] in the mesocarp was caused by a respiratory demand from seeds directly.

Increased temperature advance the onset and increases the rate of CD in Shiraz berries (Bonada *et al.*, 2013b). Using a modelling approach for pear fruit it was shown that increasing temperature should strongly increase respiration rate but not to affect the gas diffusion properties resulting in predicted very low core [O_2_] (Ho *et al.*, 2009). Our direct measures of berry mesocarp [O_2_] profiles concur with this prediction. We also observed typical Q_10_ and activation energy for respiration of 2.47 and 2.27 for whole berry respiration rates between 10 and 40 ^o^C for Chardonnay and Shiraz berries respectively, and it was only at 40 ^o^C that blocking the pedicel lenticels reduced respiration. The activation energies were similar to those reported by Hertog *et al.* (1998) for apple (52875 J/mol), chicory (67139 J/mol) and tomato (67338 J/mol). Unlike pear fruit, wine-grape berries ripe on the plant and can become considerably hotter than the surrounding air (Caravia *et al.*, 2016; Smart and Sinclair, 1976; Tarara *et al.*, 2008). Transient high temperatures would create a large respiratory demand and low [O_2_] in the centre of the mesocarp as we observed. However, subsequent cooling during the night or during milder weather will reduce the respiratory demand and result in higher internal [O_2_] if the diffusivity for O_2_ remains the same. This could then result in production of damaging ROS that may cause unrecoverable cell damage (Pfistersieber and Brandle, 1994; Rawyler *et al.*, 2002).

Finally, it is useful to consider the possible links between CD and berry dehydration. Hypoxia and anoxia are associated with reduced plasma membrane water permeability (Zhang and Tyerman, 1991) caused by closing of water channels of the plasma membrane intrinsic protein (PIP) family (Tournaire-Roux *et al.*, 2003). This is due to sensitivity to lowered cytosolic pH under hypoxia. A PIP aquaporin (*VvPIP2;1*) that is highly expressed in the ripening berry (Choat *et al.*, 2009) would be predicted to have reduced water permeation under hypoxia (Tournaire-Roux *et al.*, 2003) perhaps accounting for the decrease in whole berry hydraulic conductance that is consistently observed for Chardonnay and Shiraz (Scharwies and Tyerman, 2017; Tilbrook and Tyerman, 2009) and the decreased propensity of berries to split due to swelling in wet conditions (Clarke *et al.*, 2010). Hypoxia induced decrease in water permeability would also decrease the membrane reflection coefficient and reduce the effectiveness of the high concentrations of sugar in the mesocarp cells to oppose the tensions developed in the apoplast by the parent vine.

### Conclusion

Grape internal [O_2_] declines during fruit development and is associated with mesocarp cell death. Our data suggest the differences in O_2_ availability between cultivars could be associated with seed development and differences in lenticel morphology. 3D modelling of the air spaces in grape berries provides new insights on the pathways of O_2_ diffusion. The data presented here add to the understanding of cell death in the mesocarp of grapes late in ripening and provides a basis for further research into the role of berry gas exchange through pedicel lenticels in berry development, berry quality and cultivar selection for adapting viticulture to a warming climate.

## Supplementary Data

Table S1. Summary of berry source and traits measured

Figure S1. Temperature dependence of berry respiration rate.

Figure S2. Respiratory Q_10_ of Chardonnay and Shiraz berries in response to short-term measurement temperature at two maturity stages.

Figure S3. Micro CT analysis of air spaces Shiraz berries at two development stages.

## Acknowledgements

We thank Wendy Sullivan for expert technical assistance and Adelaide Microscopy for their facilities and technical support. This research was conducted by the Australian Research Council Training Centre for Innovative Wine Production (www.adelaide.edu.au/tc-iwp/), which is funded as a part of the ARC's Industrial Transformation Research Program (Project No IC130100005) with support from Wine Australia and industry partners.

